# Asymmetric dysregulation of glutamate dynamics across the synaptic cleft in a mouse model of Alzheimer disease

**DOI:** 10.1101/2022.04.21.489062

**Authors:** Kyle J Brymer, Emily P Hurley, Jessica C Barron, Bandhan Mukherjee, Jocelyn R Barnes, Firoozeh Nafar, Matthew P Parsons

## Abstract

Most research on glutamate spillover focuses on the deleterious consequences of postsynaptic glutamate receptor overactivation. However, two decades ago, it was noted that the glial coverage of hippocampal synapses is asymmetric: astrocytic coverage of postsynaptic sites exceeds coverage of presynaptic sites by a factor of four. The fundamental relevance of this glial asymmetry remains poorly understood. Here, we used the glutamate biosensor iGluSnFR, and restricted its expression to either CA3 or CA1 neurons to visualize glutamate dynamics at pre- and postsynaptic microenvironments, respectively. We demonstrate that inhibition of the primarily astrocytic glutamate transporter-1 (GLT-1) slows glutamate clearance to a greater extent at presynaptic compared to postsynaptic membranes. GLT-1 expression was reduced early in a mouse model of AD, resulting in slower glutamate clearance rates at presynaptic but not postsynaptic membranes that opposed presynaptic short-term plasticity. Overall, our data demonstrate that the presynapse is particularly vulnerable to GLT-1 dysfunction and may have implications for presynaptic impairments in a variety of brain diseases.

## Introduction

Tight control over the spatiotemporal dynamics of extracellular glutamate is necessary to prevent the toxic effects associated with extracellular glutamate accumulation (Danbolt, 2001). High-affinity sodium-dependent excitatory amino acid transporters (EAATs) rapidly remove glutamate from the extracellular space, thereby promoting a high signal-to-noise ratio during synaptic neurotransmission and maintaining ambient glutamate concentrations at sub-toxic levels (Lehre and Danbolt, 1998). Glutamate transporter-1 (GLT-1) is the most abundant EAAT in the brain and is expressed in both astrocytes and presynaptic terminals (Lehre and Danbolt, 1998; Rimmele and Rosenberg, 2016). Impaired glutamate clearance, notably resulting from reduced GLT-1 expression and/or function, can trigger excitotoxic cell death that is often attributed to the overactivation of postsynaptic NMDA receptors (Benarroch, 2018; Bukke et al., 2020; Esposito et al., 2013; Li and Selkoe, 2020; Parsons and Raymond, 2014; Rudy et al., 2015; Wang and Reddy, 2017). As glutamate toxicity is known to play a pivotal role in the pathogenesis of neurodegenerative diseases such as Alzheimer disease (AD), it is imperative that we increase our understanding of glutamate regulation in the healthy brain, and its dysregulation in disease.

Approximately 90-95% of the GLT-1 protein found in the brain is expressed in astrocytes (Danbolt et al., 2016; Rimmele and Rosenberg, 2016). Two decades ago, it was first observed that the glial coverage of hippocampal synapses is asymmetric; that is, glial coverage of the postsynapse exceeds that of the presynapse by a factor of four (Lehre and Rusakov, 2002). A more recent study using 3D electron microscopy confirmed that the maximum astrocyte volume fraction around postsynaptic densities was significantly larger than that surrounding presynaptic boutons (Gavrilov et al., 2018). The functional implications of this glial asymmetry — supported by modeling (Lehre and Rusakov, 2002) — suggest that glutamate spillover in the hippocampus favors the overactivation of presynaptic autoreceptors, possibly as a negative feedback mechanism to attenuate release. At present, it remains unknow whether glutamate clearance rates are differentially regulated across the synaptic cleft, and if so, how glutamate dynamics at pre- and postsynaptic microenvironments are impacted in diseases associated with GLT-1 impairment.

Impaired GLT-1 expression and/or function has been widely reported in AD animal models and in postmortem tissue (Ferrarese et al., 2000; Hefendehl et al., 2016; Li et al., 2009; Masliah et al., 1996; Mookherjee et al., 2011; Schallier et al., 2011; Scimemi et al., 2013; Scott et al., 2011; Wang and Reddy, 2017). At present, we have a poor understanding of how GLT-1 dysfunction in AD alters the spatiotemporal dynamics of glutamate at pre- and postsynaptic membranes, and therefore a poor understanding of the glutamate receptor subtypes, and the side of the synapse on which they reside, that are most heavily impacted by GLT-1 impairment. Here, we used high-speed two-photon imaging following sparse injections of the glutamate biosensor iGluSnFR targeted to CA3 or CA1 to visualize real-time glutamate dynamics at pre- or postsynaptic microenvironments, respectively, in the stratum radiatum. Our data demonstrate that at CA3-CA1 hippocampal synapses in control mice, GLT-1 inhibition slows glutamate clearance to a greater extent at presynaptic compared to postsynaptic microenvironments. In the 3xTg mouse model of AD, GLT-1 dysfunction slowed glutamate clearance at presynaptic but not postsynaptic microenvironments, resulting in presynaptic mGluR overactivation that opposed short-term plasticity. As GLT-1 dysfunction is implicated in numerous brain diseases in addition to AD (Brymer et al., 2021; Takahashi et al., 2015), our experiments may also have broader implications for presynaptic vulnerability in a range of disease states.

## Materials and Methods

### Animals

The low concentration DHK experiments presented in Fig. 1 were conducted on acute brain slices obtained from male C57BL/6NCrl mice. Mice were ordered from Charles River at ∼ four weeks of age and were acclimatized for at least three days upon arrival at Memorial University’s animal care facility. All remaining experiments were conducted on acute brain slices obtained from male and female 3xTg mice (The Jackson Laboratory strain #034830) (Oddo et al., 2003) and age-matched B6129SF2/J controls (The Jackson Laboratory strain #101045). No differences in male and female mice were noted for any of the measures obtained in the present study; therefore, the data were pooled from both sexes. All 3xTg and control mice were bred in our in-house colony at Memorial University. All mice were group housed in ventilated cage racks, provided with *ad libitum* access to standard chow and water, and maintained on a standard 12 h light/dark cycle. All experimental procedures were approved by Memorial University’s Animal Care Committee and were conducted in accordance with the guidelines set by the Canadian Council on Animal Care.

**Figure 1.**
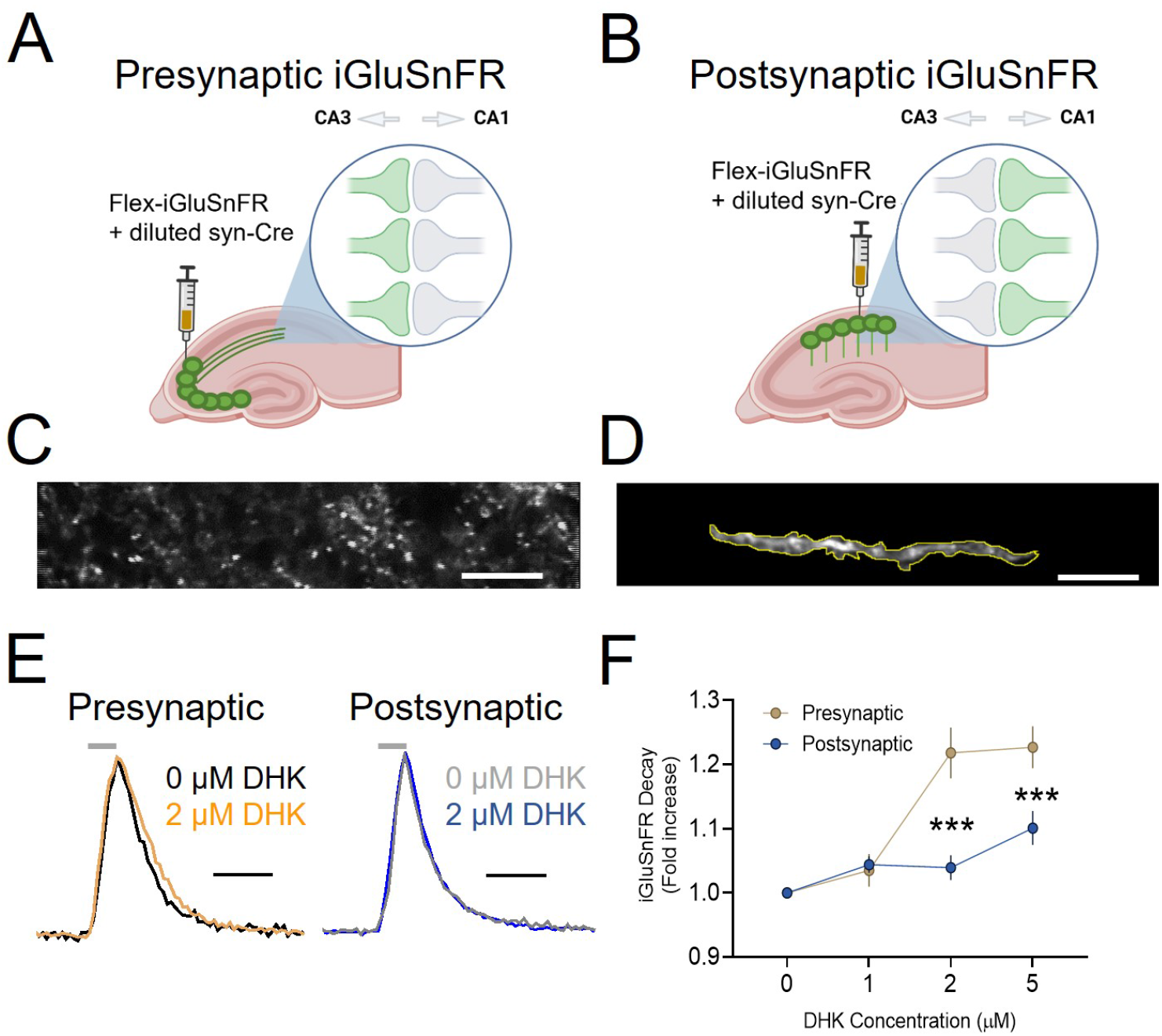
Presynaptic microenvironments are more susceptible to partial GLT-1 inhibition than postsynaptic microenvironments. Schematic showing presynaptic (A) and postsynaptic (B) expression of iGluSnFR. Two-photon microscopy images with iGluSnFR expression obtained in CA1 showing presynaptic iGluSnFR expression with punctate expression pattern (C), or postsynaptic iGluSnFR expression of a single dendritic segment (D). (E) Evoked iGluSnFR responses (5 pulses, 100 Hz) in CA1 stratum radiatum with and without DHK application. Shown are traces for pre and postsynaptic microenvironments. (F) DHK application increases iGluSnFR decay tau to a greater extent at presynaptic compared to postsynaptic membranes. Horizontal lines above iGluSnFR traces indicate the timing and duration of electrical stimulation. Scale bars in C-D: 10 µm. Scale bars in E: 100 ms. Error bars indicate s.e.m. *** p < 0.001.

### Stereotaxic Surgery

Surgical procedures were similar to previously published protocols from our lab (Barnes et al., 2020; Pinky et al., 2018). 3xTg mice, age-matched controls, and C57 mice were anesthetized by isoflurane inhalation (3%) and maintained with 1.5-2% isoflurane for the duration of the surgical procedure. Mice were secured within the ear bars of a standard stereotaxic apparatus (Stoetling), eye drops applied to lubricate the eyes, and a subcutaneous 0.9% saline injection containing 2 mg/kg meloxicam was provided to prevent dehydration during the procedure. When mice became unresponsive to a toe pinch a small amount of fur above the scalp was cut using scissors, and a 0.1 ml bolus of 0.2% lidocaine was injected below the scalp. A small incision was then made in the scalp surrounding bregma, and the underlying skull exposed. A hand drill was then used to carefully thin the skull at the desired coordinates from bregma, and a fine bent needle tip was used to peel back the last layer of skull to expose the underlying cortex while minimizing tissue damage. A Neuros 7002 Hamilton Syringe was attached to an infusion pump (Pump 11 Elite Nanomite; Harvard Apparatus), which was then secured to the stereotaxic frame. Animals used for calcium imaging received a total volume of 1 µl AAV1.Syn.Flex.GCaMP6f.WPRE.SV40 (a gift from Douglas Kim & Genie Project, Addgene plasmid # 100833, http://n2t.net/addgene:100833; RRID:Addgene_100833) injected bilaterally into the hippocampus at an injection rate of 2 nl/s. The syringe was left in place for an additional 5 min after the injection to facilitate diffusion. The syringe was slowly withdrawn, incision sutured, and mice were given a subcutaneous injection of 0.5 ml of 0.9% saline before being placed on a heating pad for ∼30 min to accelerate recovery. For glutamate imaging experiments, animals received a total of 1 µl of pAAV.hSyn.Flex.iGluSnFR.WPRE.SV40 (a gift from Loren Looger, Addgene plasmid # 98931; http://n2t.net/addgene:93931; RRID:Addgene_98931) combined with a 1:50 or 1:100 dilution of 1 ul pAAV.hSyn.cre.WPRE.hGH. The following coordinates were used with respect to bregma: CA1: 2.6 mm posterior, 2.4 mm lateral, 1.1-1.3 mm ventral to brain surface (Barnes et al., 2020); CA3: 2 mm posterior, 2.3 mm lateral, 2.3 mm ventral (Botterill et al., 2021). The classification of pre- or postsynaptic iGluSnFR expression was determined on the day of imaging when viewing the transverse acute slice under the two-photon microscope. Slices were classified as “presynaptic” when CA3 but not CA1 cell body iGluSnFR expression was present, with clear punctate iGluSnFR expression observed in CA1 stratum radiatum. Slices were classified as “postsynaptic” when CA1 cell body iGluSnFR expression was present, with clear apical dendrites observed in CA1 stratum radiatum running perpendicular to the CA1 cell body layer. For some experiments, iGluSnFR was expressed in astrocytes (using the above CA1 co-ordinates) by injecting 1 µl of pENN.AAV.GFAP.iGluSnFR.WPRE.SV40 (a gift from Loren Looger, plasmid # 98930; http.//n2t.net/addgene:98930; RRID:Addgene_98930).

### Ceftriaxone Treatment

6-month-old 3xTg and age-matched control mice were used for all ceftriaxone experiments. Similar to previous protocols from our lab (Wilkie et al., 2020, 2021), mice were treated with ceftriaxone (200 mg/kg) or sterile saline via intraperitoneal injection for 5-7 days. 24 hours after the last injection, mice were sacrificed and acute sections containing the hippocampus were obtained (described below).

### Slice Preparation

At 6 months of age (2-4 weeks after iGluSnFR injection), an age which corresponds to the emergence of an AD-like phenotype in 3xTg mice (Oddo et al., 2003), mice were anesthetized with isoflurane and decapitated. A subset of experiments were performed on mice aged to 12 months, as indicated. For the low concentration DHK experiments in Fig. 1, C57BL/6NCrl mice were sacrificed at 2 months of age. The brain was quickly removed and immersed in ice-cold oxygenated (95% O_2_/5% CO_2_) slicing solution consisting of (in mM) 125 NaCl, 2.5 KCI, 25 NaHCO_3_, 1.25 NaH_2_PO_4_, 2.5 MgCl_2_, 0.5 CaCl_2_, and 10 glucose. Slices from the brains of the 12-month-old mice were prepared in N-methyl-d-glucamine (NMDG) and HEPES solutions to improve slice health as described previously (Ting et al., 2018). Transverse hippocampal slices (350 µm) were obtained using a Precisionary compresstome. Slices were transferred to a holding chamber containing oxygenated ACSF for recovery 45 minutes before experimentation. The ACSF consisted of (in mM) 125 NaCl, 2.5 KCI, 25 NaHCO_3_, 1.25 NaH_2_PO_4_, 1 MgCl_2_, 2 CaCl_2_, and 10 glucose. Slices for electrophysiology experiments were recovered for a minimum of 90 minutes. Slices from the 12-month age group were transferred to NMDG ACSF after slicing with time-dependent sodium spiking applied exactly as described (Ting et al., 2018) and were then transferred to HEPES ACSF for an additional hour before use.

### Imaging and Image Analysis

#### Glutamate imaging

After recovery, slices were transferred to the recording chamber of a Scientifica Hyperscope, and a peristaltic pump (MP-II; Harvard Apparatus) was used to perfuse oxygenated ACSF at a flow rate of 2 ml/min. Glass stimulating electrodes were pulled using a Narishige PB-7 pipette puller to a resistance of 1-3 MΩ when filled with ACSF. Alexa Fluor 594 hydrazide sodium salt was added to the ACSF in the pipette to facilitate placement in the slice while two-photon imaging. iGluSnFR was excited using a Chameleon Vision II femtosecond pulsed laser tuned to 920 nm. iGluSnFR fluorescence was captured using a 16x objective (Nikon), and ScanImage 2019 was used to control all image acquisition parameters. The glass stimulating electrode was placed directly in the Schaffer collateral pathway within the stratum radiatum, at a depth of 50-100 µm below the slice surface, approximately 20-30 µm lateral (towards CA3) to the imaging region of interest (ROI). To visualize evoked iGluSnFR transients, fast resonant scanning was used with a 10x zoom, and the imaging field was collapsed to 100 lines to achieve a frame rate of 153.6 frames per second. Clampex software (Molecular Devices) was used to send TTL triggers through the digital outputs of a Digidata 1550A (Molecular Devices) to trigger image acquisition and electrical stimulation (100 µs pulses, 75 µA) via an Iso-Flex stimulus isolator (AMPI). Five evoked iGluSnFR responses (20 second intervals) were averaged for each stimulation protocol. Values from the average file obtained from trials with presynaptic iGluSnFR expression were converted to Δ%*F*/*F*, and decay tau values were calculated in GraphPad Prism 9. Decay tau values were quantified starting at the offset of electrical stimulation (i.e. from the glutamate response at the end of the 5 or 100 pulse train to its return to baseline), similar to that described previously for widefield iGluSnFR transients (Barnes et al., 2020; Pinky et al., 2018; Wilkie et al., 2020, 2021). In slices with postsynaptic iGluSnFR expression, Δ%*F*/*F* values were obtained from a ROI drawn around a segment of an individual dendrite.

#### Calcium imaging

After recovery, slices were transferred to the recording chamber, and a peristaltic pump (MP-II; Harvard Apparatus) was used to perfuse oxygenated ACSF at a flow rate of 2 ml/min. Glass stimulating electrodes were pulled using a Narishige PB-7 pipette puller to a resistance of 1-3 MΩ when filled with ACSF. The stimulating electrode was placed directly in the Schaffer collateral pathway within the stratum radiatum, at a depth of 50-100 µm below the slice surface. Clampex software (Molecular Devices) was used to send TTL triggers through the digital outputs of a Digidata 1550A (Molecular Devices) for precise control over a LED illumination source (Prior, Lumen 300), an EM-CCD camera (Andor, iXon Ultra 897), and an Iso-flex stimulus isolator (AMPI). GCaMP6f responses to evoked neural activity were recorded with Andor Solis software, using 4 × 4 binning and an acquisition rate of 205 frames per second. Evoked GCaMP6f responses were quantified within a 10 × 10 pixel ROI (1 pixel at 4 × 4 binning = 16 µm) placed adjacent to the location of the stimulating electrode and converted to %Δ*F/F*. The area under the curve (AUC) was calculated before and after bath application of D-APV (50 µM) to determine how much of the pre- or postsynaptic calcium response was NMDA receptor-dependent.

### Electrophysiology

Acute hippocampal slices from 6-month-old 3xTg and age-matched control mice were placed in the recording chamber and were left for a minimum of 10 minutes before electrode placement. ACSF was superfused into the recording chamber at a rate of 2 ml/min. A glass stimulating electrode filled with ACSF (1-3 MΩ) was placed in the stratum radiatum to stimulate Schaffer collateral fibres in the CA1 region. A glass recording electrode filled with ACSF (1-3 MΩ) was next placed ∼400 µm from the stimulating electrode, and signals were amplified and low-pass filtered at 10 kHz with a Multiclamp 700B amplifier (Molecular Devices). Using an inter-pulse interval of 50 ms, we measured paired-pulse ratios (PPR) by dividing the fEPSP amplitude evoked by the second stimulus by the that induced by the first. We first recorded PPR for 3 minutes to establish a baseline, then applied HFS (3 × 100 Hz, 1 s, 10 second inter-train intervals). After HFS, PPR was again recorded for 5 minutes. All data were collected and analyzed using pClamp 10 software (Molecular Devices).

### Pharmacology

All drugs used in the current experiments were from Tocris Bioscience. Drugs used in the study and their concentrations are as follows: dihydrokainic acid (DHK), a competitive and selective EAAT2 blocker (EAAT2-2; 300 µM; low DHK experiments used 1, 2, or 5 µM); _DL_-threo-β-benzloxsapartic acid (_DL_-TBOA), a competitive and nonselective EAAT blocker (100 µM); DNQX disodium salt, an AMPA/kainite receptor antagonist (20 µM); _D_-AP-5, a selective NMDA receptor antagonist (50 µM); MSOP, a selective group III metabotropic glutamate receptor antagonist (100 µM); and MTEP hydrochloride, a selective group 5 metabotropic glutamate receptor antagonist (100 µM). In DHK, TBOA, or MSOP/MTEP experiments, slices were incubated for 5-10 minutes before imaging or electrophysiology was conducted.

### GLT-1 Immunohistochemistry

Mice were perfused and brains were cryoprotected in 30% sucrose until sunk. Whole brains were rapidly frozen and sliced at 20 µm on a cryostat (Leica CM3050 S), and sections were mounted directly onto gelatin-coated glass slides and stored at -80ºC until use. Day 1: Slides were removed from -80ºC and brought to room temperature, and a Dako pen was used to trace the perimeter of the slide. Slides were washed three times in 0.01 M PBS for 10 minutes each. Slides were then incubated in blocking serum (0.01 M PSB with 5% BSA + 0.2% Triton-X) for one hour. After blocking, slides were incubated in primary antibody (Guinea pig anti-GLT1, 1:500, Abcam, AB1783) overnight at 4ºC. Slides were then washed three times in 0.01 M PBS for 10 minutes each and incubated in secondary antibody (Alexa Flour 647-conjugated Donkey anti-Guinea pig, 1:500, Jackson Immunoresearch Labs) at room temperature for 2 hours. After the final washes in 3 times in 0.01 M PBS for 10 minutes each, slides were cover slipped using Dako mounting medium containing DAPI. Images of CA1 stratum radiatum were obtained at 20x on a Zeiss Axio Observer. GLT-1 intensity was calculated within an ROI in the stratum radiatum using ImageJ. For GLT-1 immunohistochemistry and imaging, all samples were processed at the same time and imaging parameters, including LED intensity and exposure times, were kept constant.

### Experimental design and statistics

Statistical tests used included unpaired t-tests and two-way repeated-measures (RM) ANOVA with Bonferroni post-hoc tests. The statistical test used for each experiment is indicated in the results text. *P* values of <0.05 were considered significant. Where indicated, *N* and *n* refer to the number of animals and slices used in each experiment, respectively.

## Results

### The presynapse is more sensitive than the postsynapse to GLT-1 inhibition

The previously observed asymmetric glial coverage across the synapse (Gavrilov et al., 2018; Lehre and Rusakov, 2002) suggests that presynaptic membranes may be more susceptible to glutamate spillover than postsynaptic membranes. To test for functional evidence of a presynaptic vulnerability to glutamate spillover during partial GLT-1 impairment, we examined the effects of sub-saturating concentrations (1-5 µM, saturating = 300 µM (Pinky et al., 2018)) of the GLT-1 inhibitor DHK on real-time glutamate dynamics sensed at presynaptic (Fig. 1A, C) or postsynaptic (Fig. 1B, D) membranes using sparse injections of iGluSnFR targeted to CA3 or CA1, respectively. These experiments were performed on healthy control (C57) mice. Both D-APV (50 µM) and DNQX (20 µM) were added to the bath solution, and two-photon microscopy was used to capture synaptically-evoked iGluSnFR transients in acute hippocampal slices. Clear iGluSnFR transients (Fig. 1E) were evoked by 5 pulses of electrical stimulation (100 Hz; 75 µA) applied to the Schaffer collaterals, and we monitored the glutamate response in baseline conditions and then to increasing sub-saturating concentrations of DHK. The decay tau of evoked iGluSnFR transients were used to quantify relative changes in the rate of glutamate clearance as described previously (Armbruster et al., 2016; Brymer et al., 2021; Pinky et al., 2018). iGluSnFR decay was more sensitive to partial GLT-1 inhibition when iGluSnFR was expressed presynaptically compared to postsynaptically (Fig. 1F; presynaptic N = 3, n = 5; postsynaptic N = 3, n = 8; RM two-way ANOVA: p_DHK_ < 0.001; *p*_location_ = 0.004; *p*_interaction_ < 0.001;). Specifically, 2 µM and 5 µM DHK significantly slowed presynaptic iGluSnFR decay to a greater extent than postsynaptic iGluSnFR (Bonferroni p < .001 for both 2 µM and 5 µM). Thus, at CA3-CA1 synapses in control mice, presynaptic membranes appear to be more sensitive to the effects of partial GLT-1 inhibition, in agreement with the morphological asymmetry of glial membranes favoring postsynaptic protection at the expense of the presynapse (Gavrilov et al., 2018; Lehre and Rusakov, 2002). As a positive control to help confirm our ability to separate pre- and postsynaptic biosensor responses with our AAV injections, we injected WT mice (2-4 months) with the calcium indicator GCaMP6f into either CA3 (for presynaptic expression) or CA1 for postsynaptic expression. Calcium responses in the stratum radiatum were induced by high frequency stimulation (HFS; 100 pulses, 100 Hz) applied to the Schaffer collaterals; this stimulation protocol is commonly used to evoke long-term potentiation (LTP) that is dependent upon calcium influx through postsynaptic NMDA receptors (NMDARs) (Malenka and Bear, 2004; Nicoll, 2017). We reasoned that NMDAR blockade with D-APV should significantly block the postsynaptic calcium response to HFS, while having minimal or no effect on presynaptic calcium responses. Indeed, postsynaptic GCaMP6f responses to HFS were significantly inhibited by NMDAR antagonism, while presynaptic GCaMP6f responses were completely unaffected by D-APV (Supplementary Fig. 1A-C; presynaptic n = 6, postsynaptic n = 11; t-test: p < .001;). These data demonstrate that our AAV injections can effectively isolate pre- and postsynaptic membrane responses to evoked activity.

### Impaired glutamate clearance at presynaptic microenvironments in 3xTg mice

Our low concentration DHK experiments above suggest partial GLT-1 inhibition slows glutamate clearance to a greater extent at presynaptic compared to postsynaptic microenvironments. GLT-1 function and/or expression is partially decreased in a variety of neurodegenerative diseases, including AD (Brymer et al., 2021). Therefore, we hypothesized that a similar presynaptic vulnerability to glutamate uptake deficits may be apparent in 3xTg mice, a commonly-used mouse model of AD that presents with age-dependent amyloid and tau pathology (Oddo et al., 2003). Acute slices were obtained from mice at six months of age, and neural activity was evoked with a short burst (5 pulses at 100 Hz) as well as with a longer train (100 pulses at 100 Hz) of electrical stimulation, the latter being known to overwhelm the glutamate uptake system (Diamond and Jahr, 2000; Pinky et al., 2018). Despite significantly reduced GLT-1 expression in 3xTg mice at this age (Supplementary Fig. 2; WT n = 12, 3xTg n = 10; t-test: p = 0.002), postsynaptic iGluSnFR decay did not differ between the genotypes, even when challenged with the longer train of neural activity (Fig. 2A-D; WT N=7, n=14; 3xTg N=6, n=14; RM two-way ANOVA: p_pulsenumber_ < 0.001; p_genotype_ = 0.285; p_interaction_ = 0.602). Postsynaptic iGluSnFR response peaks also did not differ between the two genotypes (Supplementary Fig. 3A; *p*_pulsenumber_ < 0.001; *p*_genotype_ =.418; *p*_interaction_ =.418).

**Figure 2.**
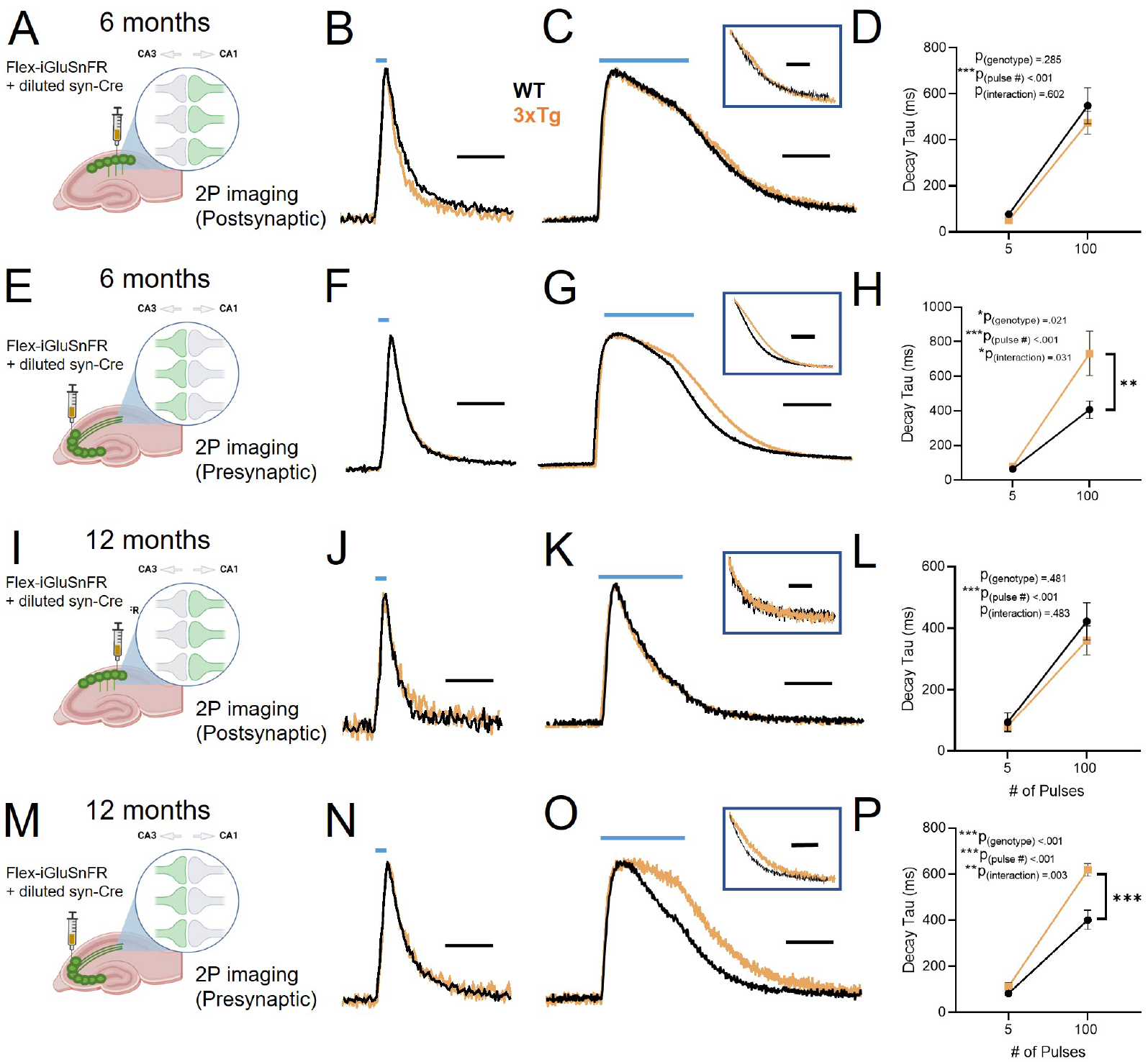
Slow glutamate clearance at pre- but not postsynaptic microenvironments in the 3xTg hippocampus. (A-D) Postsynaptic iGluSnFR expression in CA1 dendrites (A). Average iGluSnFR responses to 5 pulses (B) and 100 pulses (C) in six-month-old WT (black) and 3xTg (orange) mice. Grouped data shown in D. (E-H) Presynaptic iGluSnFR expression in CA1 dendrites (E). Average iGluSnFR responses to 5 pulses (F) and 100 pulses (G) in six-month-old WT (black) and 3xTg (orange) mice. Grouped data shown in H. (I-P) Same as A-H but in 12-month-old WT and 3xTg mice. Boxed traces in C, G, K, O depict average iGluSnFR traces normalized to the iGluSnFR value at the end of the one-second stimulation. Blue lines above iGluSnFR traces indicate the timing and duration of electrical stimulation. Scale bars in B, F, J, N: 200 ms. Scale bars in C, G, K, O: 500 ms. Error bars indicate s.e.m. ** p < 0.01, *** p < 0.001.

In contrast, presynaptic iGluSnFR decay was significantly slower in 3xTg compared to WT mice, specifically when challenged with the longer train of activity (Fig. 2E-H; WT N = 6, n = 12; 3xTg N = 6, n = 11; RM two-way ANOVA: p_pulsenumber_ < 0.001; p_genotype_ = 0.021; p_interaction_ = 0.031). At 5 pulses of stimulation, 3xTg and WT mice did not differ in time taken to clear extracellular glutamate (Bonferroni p = 0.999) but increasing the number of pulses to 100 revealed a clear deficit in glutamate clearance in 3xTg mice (Bonferroni p = 0.003), demonstrating that the deficit is only revealed when the glutamate transporter system is challenged with a long duration of activity. The maximum amount of glutamate released was not different between the two genotypes, as the peak responses of presynaptic iGluSnFR expression was similar between WT and 3xTg mice (Supplementary Fig. 3B; RM two-way ANOVA: p_pulsenumber_ < 0.001; p_genotype_ = 0.289; p_interaction_ = 0.949). The observed impairment of glutamate clearance at presynaptic but not postsynaptic membranes in 3xTg mice was also observed in a separate cohort of mice where the ACSF was heated to 32 ºC (Supplementary Fig. 4; Postsynaptic: WT N = 3, n = 9; 3xTg N = 3, n = 9; RM two-way ANOVA: p_pulsenumber_ < 0.001; p_genotype_ = 0.500; p_interaction_ = 0.353; Presynaptic: WT N = 3, n = 8; 3xTg N = 4, n = 10; RM two-way ANOVA: p_pulsenumber_ < 0.001; p_genotype_ = 0.030; p_interaction_ = 0.026).

The above experiments were conducted in six-month-old mice. In the 3xTg AD model, this age corresponds to the early emergence of an AD-like phenotype, including mild spatial-memory impairment and deficits in synaptic transmission (Oddo et al., 2003; Stover et al., 2015). Next, we asked if our observed asymmetric dysregulation of glutamate clearance across the synaptic cleft persisted to a later disease stage, or whether more global impairments (i.e. at both pre- and postsynaptic microenvironments) could be observed. At 12 months of age, 3xTg mice continued to display similar iGluSnFR responses as control mice when iGluSnFR was expressed in CA1 dendrites (Fig. 2I-L; WT N = 5, n = 10; 3xTg N = 5, n = 12; RM two-way ANOVA: p_pulsenumber_ < 0.001, p_genotype_ = 0.481, p_interaction_ = 0.483). Similar to our findings at six months, when expressed presynaptically, iGluSnFR decay tau values were significantly longer in 3xTg mice compared to WT (Fig. 2M-P; WT N = 4, n = 8; 3xTg N = 4, n = 12; RM two-way ANOVA: p_pulsenumber_ < 0.001, p_genotype_ < 0.001, p_interaction_ = 0.003). Again, glutamate clearance was significantly slower in 3xTg mice following 100 pulses (Bonferroni p < 0.001) but not 5 pulses (Bonferroni p = 0.380), suggesting that glutamate transporters need to be overwhelmed (Diamond and Jahr, 2000; Pinky et al., 2018) to observe a clear genotype difference. Together, these data demonstrate relative postsynaptic protection and presynaptic vulnerability to glutamate uptake impairments in both early and later disease stages in 3xTg mice. All remaining experiments were conduced on mice six months of age.

### GLT-1 dysfunction underlies the slow glutamate clearance at 3xTg presynaptic microenvironments

Glutamate uptake in the hippocampus is mediated by GLT-1, GLAST and, to a lesser extent, EAAC1 (Brymer et al., 2021; Danbolt, 2001). To determine whether the slow glutamate clearance observed at presynaptic membranes in 3xTg mice was due specifically to GLT-1 dysfunction, we probed the contribution of GLT-1 to overall glutamate clearance rates by bath applying a saturating concentration of DHK (300 µM (Pinky et al., 2018)), applied in the presence of 50 µM D-APV and 20 µM DNQX). We then quantified the GLT-1 decay ratio, calculated by the decay tau fold increase induced by DHK; a larger decay ratio being indicative of a larger contribution of GLT-1 in the overall glutamate clearance rate (Hanson et al., 2015; Pinky et al., 2018). When we applied DHK to slices with presynaptic iGluSnFR expression, we found that the GLT-1 decay ratio was significantly reduced in 3xTg mice (Fig. 3A-B; WT N = 5, n = 10; 3xTg N = 3, n = 11; RM two-way ANOVA: p_pulsenumber_ < 0.001; p_genotype_ = 0.004; p_interaction_ = 0.093). Peak iGluSnFR responses were not significantly different in the two genotypes following DHK application (Supplementary Fig. 5A; RM two-way ANOVA: p_pulsenumber_ < 0.001; p_genotype_ = 0.320; p_interaction_ = 0.029). Next, we asked whether GLT-1 upregulation via ceftriaxone could reverse the glutamate clearance deficit seen in 3xTg mice. We, like many others, have shown that 5-7 consecutive days of ceftriaxone (200 mg/kg, i.p.) administration increases GLT-1 in the hippocampus (Wilkie et al., 2021). Saline-treated mice showed the same effect as we had observed previously; namely, that glutamate clearance at presynaptic microenvironments is significantly slower in 3xTg mice. However, in these slices with presynaptic iGluSnFR expression, ceftriaxone completely restored 3xTg glutamate clearance rates to WT levels (Fig. 3C-D WT saline N = 3, n = 8; 3xTg saline N = 5, n = 11; WT ceftriaxone N = 3, n = 9; 3xTg ceftriaxone N = 3, n = 10; RM two-way ANOVA: p_pulsenumber_ < 0.001; p_genotype_ < 0.001; p_interaction_ = 0.008). Together, these results demonstrate that the GLT-1 impairment observed in 3xTg mice results in prolonged glutamate exposure to presynaptic membranes in the stratum radiatum.

**Figure 3.**
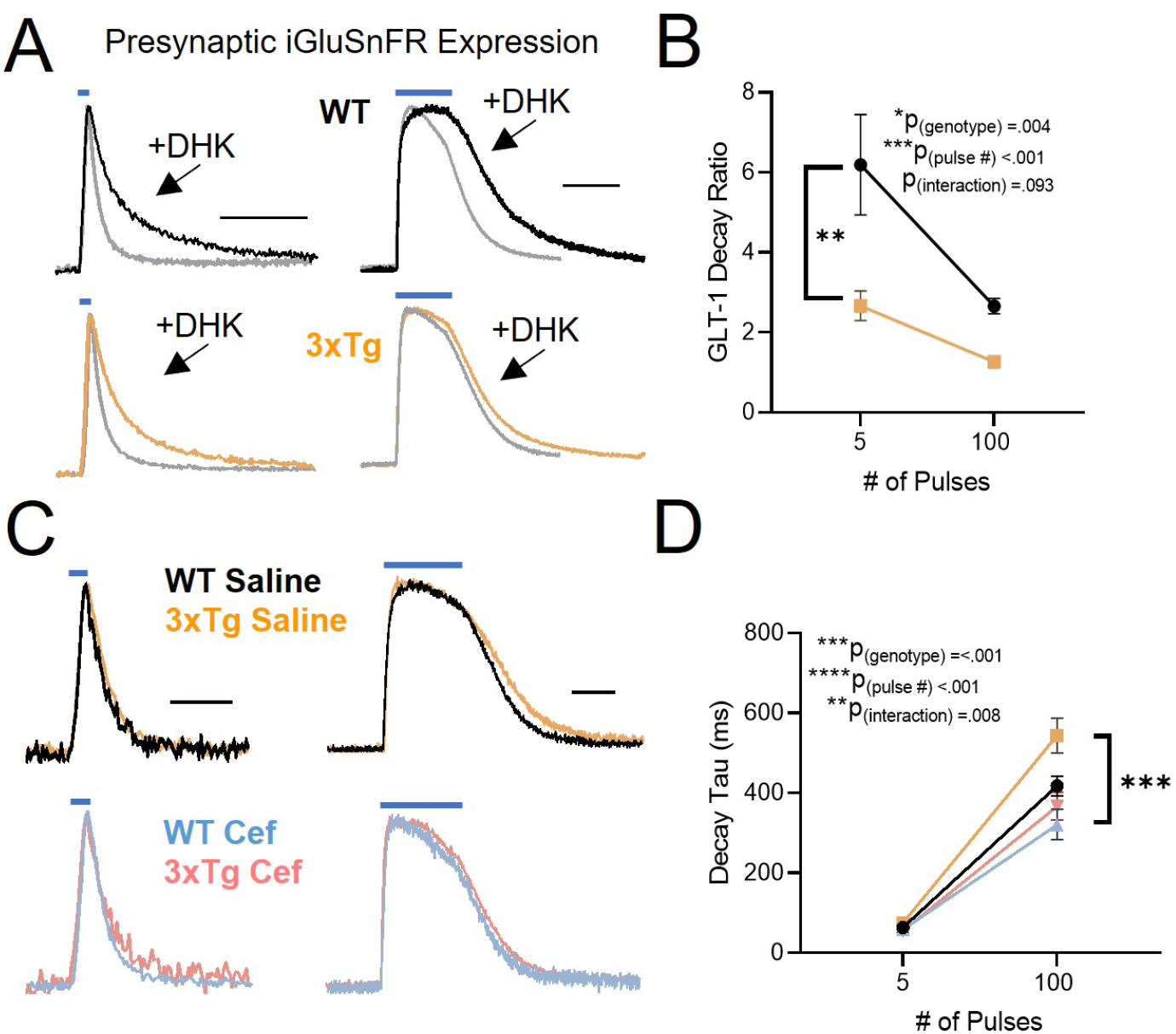
GLT-1 is dysfunctional in presynaptic microenvironments of the 3xTg hippocampus. (A) Average presynaptic iGluSnFR response to 5 (left) and 100 pulses (right) of stimulation (100 Hz) in WT (top; black and grey) and 3xTg mice (bottom; orange and grey) before and after DHK application. Grey traces denote average iGluSnFR response before DHK application. (B) GLT-1 decay ratio in WT and 3xTg mice, calculated by the fold increase in iGluSnFR decay induced by a saturating concentration of the GLT-1 inhibitor DHK. (C) Average postsynaptic iGluSnFR responses to 5 (left) and 100 pulses (right) of stimulation in WT-saline (black), 3xTg-saline (orange), WT-ceftriaxone (Cef; blue), and 3xTg-ceftriaxone-treated mice (pink). (D) iGluSnFR decay tau in saline- or ceftriaxone-treated WT and 3xTg mice. Blue lines above iGluSnFR traces indicate the timing and duration of electrical stimulation. Scale bars in A: 500 ms (left) and 1000 ms (right). Scale bars in C: 200 ms (left) and 500 ms (right). Error bars indicate s.e.m. ** p < 0.01, *** p < 0.001.

Next, we repeated the same DHK and ceftriaxone experiments, but for postsynaptic iGluSnFR responses isolated in individual CA1 dendrites. In stark contrast to presynaptic iGluSnFR, we found that that the postsynaptic GLT-1 decay ratio was enhanced, rather than reduced, in 3xTg mice (Fig. 4A-B; WT N = 3, n = 9; 3xTg N = 4, n = 11; RM two-way ANOVA: p_pulsenumber_ < 0.001; p_genotype_ = 0.011; p_interaction_ = 0.015). Peak glutamate levels remained unchanged between the genotypes in the presence of DHK (Supplementary Fig. 5B; RM two-way ANOVA: p_pulsenumber_ < 0.001; p_genotype_ = 0.087; p_interaction_ = 0.379). Again, in contrast to presynaptic iGluSnFR expression, ceftriaxone had no significant effect on postsynaptic iGluSnFR dynamics (Fig. 4C-D; WT saline *N* = 5, *n* = 8; 3xTg saline *N* = 5, *n* = 8; WT ceftriaxone *N* = 3, *n* = 10; 3xTg ceftriaxone *N* = 3, *n* = 8; RM two-way ANOVA: *p*_pulsenumber_ < 0.001; *p*_genotype_ = 0.157; *p*_interaction_ = 0.175). Together, these results suggest that GLT-1 can have both a reduced (presynaptic) and enhanced (postsynaptic) contribution to overall glutamate clearance, depending upon the microenvironment under study. Furthermore, our data suggest that GLT-1 upregulation may be beneficial in 3xTg mice by restoring presynaptic glutamate dynamics to WT levels.

**Figure 4.**
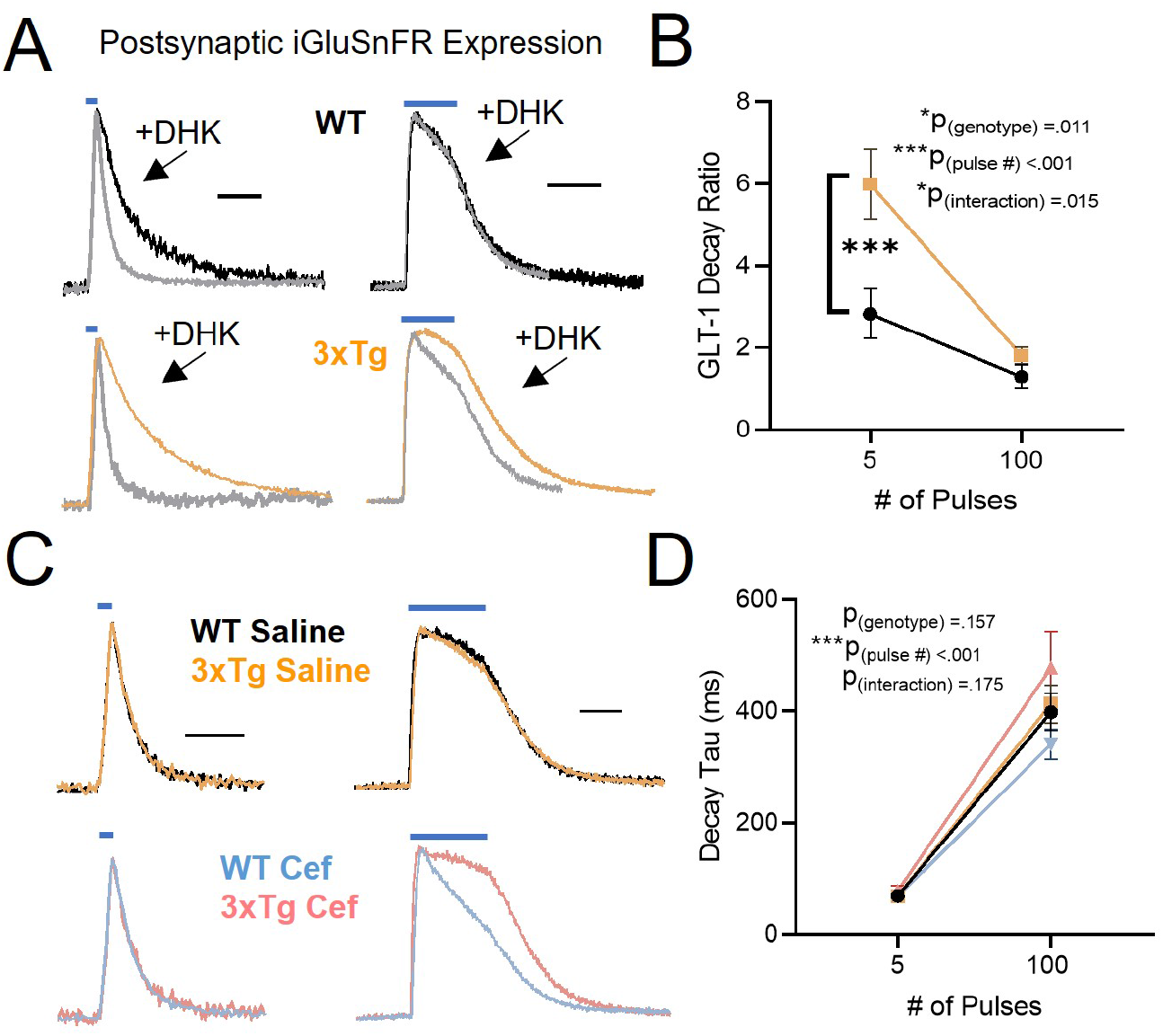
GLT-1 manipulation does not dramatically alter glutamate dynamics in postsynaptic microenvironments of the 3xTg hippocampus. (A) Average postsynaptic iGluSnFR response to 5 (left) and 100 pulses (right) of stimulation (100 Hz) in WT (top; black and grey) and 3xTg mice (bottom; orange and grey) before and after DHK application. Grey traces denote average iGluSnFR response before DHK application. (B) GLT-1 decay ratio in WT and 3xTg mice, calculated by the fold increase in iGluSnFR decay induced by a saturating concentration of the GLT-1 inhibitor DHK. (C) Average postsynaptic iGluSnFR responses to 5 (left) and 100 pulses (right) of stimulation in WT-saline (black), 3xTg saline (orange), WT-ceftriaxone (Cef; blue), and 3xTg-ceftriaxone-treated mice (pink). (D) GLT-1 decay ratio in saline- or ceftriaxone-treated WT and 3xTg mice. Blue lines above iGluSnFR traces indicate the timing and duration of electrical stimulation. Scale bars in A: 500 ms (left) and 1000 ms (right). Scale bars in C: 200 ms (left) and 500 ms (right). Error bars indicate s.e.m. *** p < 0.001.

Our DHK and ceftriaxone experiments suggested that GLT-1 impairments mediate the slow clearance rates observed at 3xTg presynaptic membranes. Nonetheless, we asked whether glutamate diffusion could differ between the genotypes and/or the pre- and postsynaptic microenvironments of interest in the present study. To this end, we applied a saturating concentration of the non-selective glutamate transporter blocker DL-TBOA (100 µM, applied in the presence of 50 µM D-APV and 20 µM DNQX). Under these conditions, iGluSnFR decay values are dramatically slowed compared to control levels, and primarily reflect the diffusion of glutamate away from the imaging ROI (Pinky et al., 2018). We found that relative glutamate diffusion rates, as estimated by iGluSnFR decay tau in the presence of saturating TBOA, were not significantly different between genotypes regardless of whether iGluSnFR expression was presynaptic (Supplementary Fig. 6A-C; WT *N* = 4, *n* = 8; 3xTg *N* = 6, *n* = 8; RM two-way ANOVA: *p*_pulsenumber_ < 0.001; *p*_genotype_ = 0.650; *p*_interaction_ = 0.058) or postsynaptic (Supplementary Fig. 6D-F; WT *N* = 4, *n* = 10; 3xTg *N* = 5, *n* = 9; RM two-way ANOVA: *p*_pulsenumber_ < 0.001; *p*_genotype_ = 0.263; *p*_interaction_ = 0.810). Similarly, iGluSnFR peak values in the presence of TBOA were not significantly different between genotypes for either presynaptic (Supplementary Fig. 7A; RM two-way ANOVA: *p*_pulsenumber_ = 0.010; *p*_genotype_ = 0.808; *p*_interaction_ = 0.865) or postsynaptic expression (Supplementary Fig. 7B; RM two-way ANOVA: *p*_pulsenumber_ < 0.001; *p*_genotype_ = 0.422; *p*_interaction_ = 0.232). Together with our previous results, the data suggest that poor GLT-1-mediated uptake, not poor diffusion, is primarily responsible for the slow glutamate clearance rates observed at 3xTg presynaptic membranes.

### Slow glutamate clearance is also observed at glial membranes in 3xTg mice

Our data suggest that GLT-1 dysfunction slows glutamate clearance primarily at presynaptic membranes in 3xTg mice. In contrast, glutamate clearance rates at postsynaptic membranes were similar between WT and 3xTg mice, consistent with the asymmetric glial coverage of tripartite synapses that enhances postsynaptic protection from spillover (Lehre and Rusakov, 2002). Next, we asked whether the observed uptake impairment exclusively impacted presynaptic membranes, or whether a more general approach to assess extrasynaptic glutamate levels could also detect slower clearance in 3xTg mice. Determining the precise microenvironments most susceptible to glutamate uptake impairments is of interest as glutamate receptors are located on glial cells in addition to their more canonical pre- and postsynaptic locations (Lalo et al., 2021; Skowrońska et al., 2019). We injected six-month-old WT and 3xTg mice with iGluSnFR as before, but this time iGluSnFR expression was restricted to astrocytes through the use of the GFAP promoter (Marvin et al., 2013). Similar to our results for presynaptic iGluSnFR expression, glutamate clearance rates were slower in 3xTg mice compared to WT (Supplementary Fig. 8; WT N = 3, n = 12; 3xTg N = 3, n = 10; RM two-way ANOVA: p_pulsenumber_ < 0.001; p_genotype_ = 0.004; p_interaction_ = 0.006) with 3xTg mice being significantly slower at clearing glutamate at 100 pulses (Bonferroni: p <.001). These results suggest that the slow glutamate clearance from the extracellular space in 3xTg mice primarily affects presynaptic and astrocytic membranes over postsynaptic dendrites.

### Ceftriaxone or mGluR antagonists prevent short-term presynaptic plasticity deficits in the 3xTg hippocampus

The prolonged glutamate transients observed at presynaptic membranes in 3xTg mice suggest that GLT-1 dysfunction could promote presynaptic autoreceptor overactivation. To test this hypothesis, we examined post-tetanic potentiation (PTP), a form of short-term plasticity characterized by a temporary facilitation of presynaptic release (Zucker and Regehr, 2002). PTP is impaired in 3xTg mice (Chakroborty et al., 2019), though the underlying mechanisms of the impairment are poorly understood. In acute hippocampal sections obtained from six-month-old WT and 3xTg mice, we found that PTP, induced by HFS (100 pulses, 1 s), was significantly impaired in 3xTg mice compared to WT (Fig. 5A-B; WT N = 6, n = 15; 3xTg N = 8, n = 16; t-test: p = 0.009). In WT slices, PTP was associated with an increase in release probability, as demonstrated by a reduced PPR immediately following HFS. The magnitude of the HFS-induced change in release probability was significantly reduced in 3xTg compared to WT mice (Fig. 5C-D; WT N = 6, n = 15; 3xTg N = 8, n = 16; t-test: p = 0.031). Next, we asked whether the release facilitation associated with PTP was opposed by glutamate autoreceptor overactivation in 3xTg mice. Presynaptic glutamate receptors known to exist at adult CA3-CA1 synapses include mGluR5 and mGluR7, both of which have been shown to negatively regulate glutamate release (Gereau IV and Conn, 1995; He et al., 2019; Klar et al., 2015; Pittaluga, 2016; Shigemoto et al., 1997). We found that PTP was no longer impaired in 3xTg mice compared to WT when we blocked mGluR5 (Fig. 5E-F; WT N = 5, n = 8; 3xTg N = 5, n = 10; t-test: p = 0.888) or mGluR7 (Fig. 5I-J; WT N = 5, n = 10; 3xTg N = 5, n = 11; t-test: p = 0.434) with MTEP or MSOP, respectively. In the presence of MTEP (Fig. 5G-H; t-test: p = 0.667) or MSOP (Fig. 5K-L; t-test: p = 0.442), HFS had a similar effect on release probability in WT and 3xTg mice. These data demonstrate that blockade of either mGluR5 or mGluR7 prevents the PTP impairment in 3xTg mice. As both mGluR5 and mGluR7 have been shown to be present presynaptically at CA3-CA1 synapses where they can negatively regulate glutamate release, we reasoned that the slow glutamate clearance may be required to promote their overactivation to oppose PTP. We have shown that ceftriaxone prevents the prolonged glutamate transients at 3xTg terminals; therefore, we also assessed PTP in ceftriaxone-treated mice in the absence of mGluR5 or mGluR7 blockade. Ceftriaxone alone was sufficient to prevent the PTP impairment in 3xTg mice (Fig. 5M-N; WT N = 5, n = 13; 3xTg N = 5, n = 17; t-test: p = 0.995). Analysis of PPR demonstrated that HFS had a similar effect on release probability in both WT and 3xTg mice after ceftriaxone treatment (Fig. 5O-P; t-test: p = 0.225). Overall, our data suggest that GLT-1 dysfunction prolongs glutamate actions at the presynapse in 3xTg mice, promoting autoreceptor-mediated opposition to presynaptic short-term plasticity. Blocking either mGluR5, mGluR7 or enhancing GLT-1 with ceftriaxone is sufficient to restore short-term plasticity to control levels.

**Figure 5.**
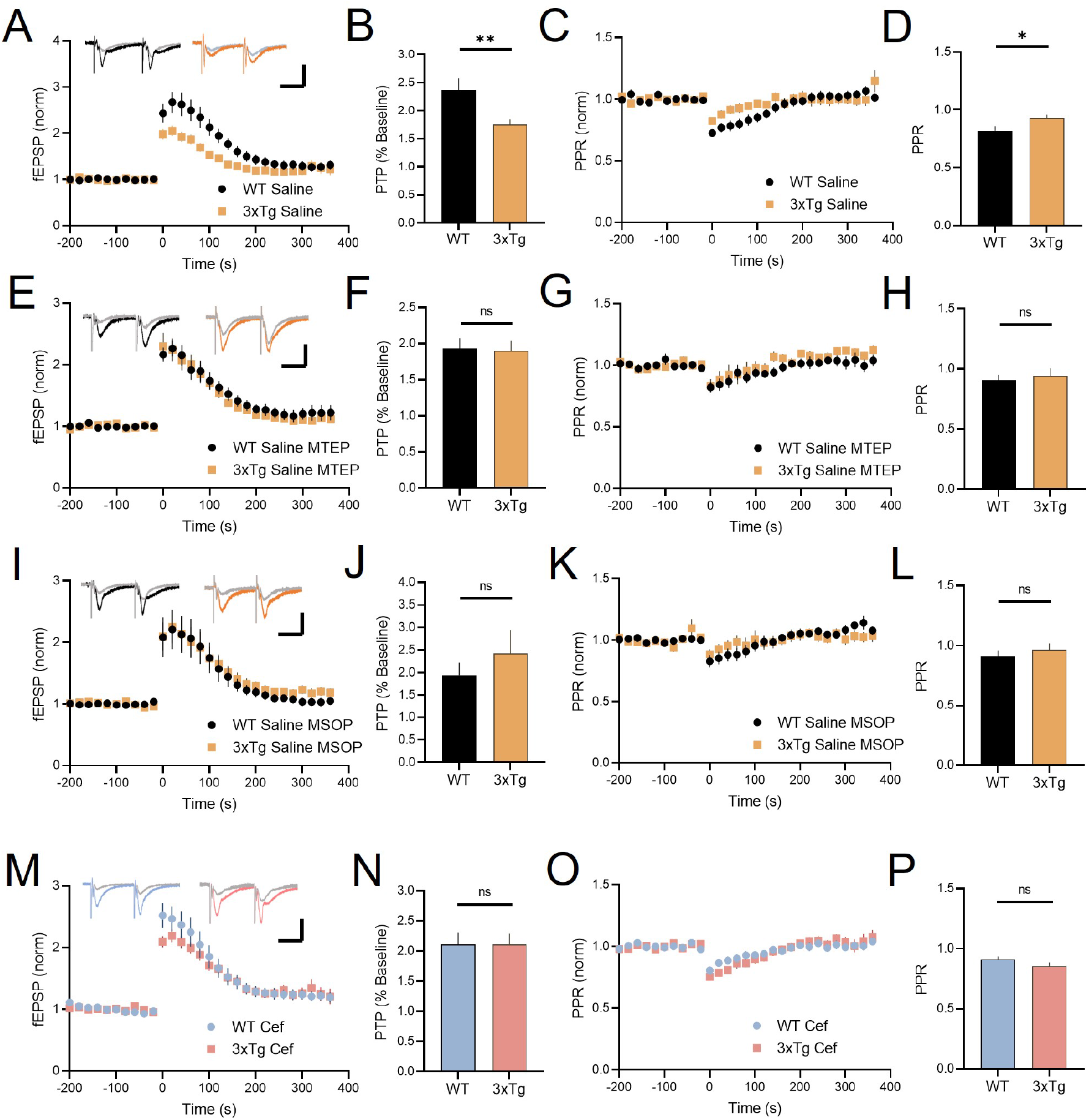
mGluR antagonism or ceftriaxone is sufficient to prevent short-term plasticity impairment in 3xTg mice. (A) Post-tetanic potentiation (PTP) in WT (black) and 3xTg (orange). PTP is induced by high-frequency stimulation (HFS) at time = 0. (B) PTP is significantly decreased in 3xTg mice. (C) Paired-pulse ratio (PPR) in WT and 3xTg. (D) PPR changes after HFS are impaired in 3xTg mice. (E-H) PTP and PPR responses to HFS in WT and 3xTg mice during bath application of MTEP (100 µM). (I-L) PTP and PPR responses to HFS in WT and 3xTg mice during bath application of MSOP (100 µM). (M-P) PTP and PPR responses to HFS in WT and 3xTg mice treated with ceftriaxone. Scale bars: 25 ms, 500 µV. Error bars indicate s.e.m. *p < 0.05, ** p < 0.01. ns, not significant.

## Discussion

Here, we demonstrate that compromised GLT-1-mediated uptake, either through pharmacological inhibition or in the 3xTg mouse model of AD, slows glutamate clearance to a greater extent at presynaptic compared to postsynaptic membranes in the hippocampus. The asymmetric uptake deficit was demonstrated in multiple experimental 3xTg cohorts, including mice at different disease stages (6 and 12 months), different bath temperatures, and in saline- (but not ceftriaxone-) treated mice. Based on the asymmetrical glial coverage of tripartite synapses in the stratum radiatum, it was previously proposed that glutamate spillover should have a greater impact at the presynapse than the postsynapse (Gavrilov et al., 2018; Lehre and Rusakov, 2002). By visualizing glutamate dynamics independently in CA3 axons or CA1 dendrites, we provide strong functional data supporting such a presynaptic vulnerability to glutamate uptake impairments. Subsequent electrophysiological experiments suggest that the presynaptic vulnerability to GLT-1 impairment in 3xTg mice promotes autoreceptor-mediated opposition to short-term presynaptic plasticity.

### Presynaptic vulnerability to GLT-1 dysfunction in 3xTg mice

GLT-1 reductions are widely reported in AD brains of human patients and animal models (Ferrarese et al., 2000; Hefendehl et al., 2016; Li et al., 2009; Masliah et al., 1996; Mookherjee et al., 2011; Schallier et al., 2011; Scimemi et al., 2013; Scott et al., 2011; Wang and Reddy, 2017). We also observed reduced GLT-1 immunofluorescence in CA1 stratum radiatum of 3xTg mice in the present study. A saturating concentration of DHK had a smaller effect on iGluSnFR decay in 3xTg mice than it did in WT mice, but only when iGluSnFR was expressed presynaptically. Therefore, GLT-1 appears to play a reduced role in glutamate clearance from presynaptic microenvironments in 3xTg tissue. It is conceivable that such presynaptic vulnerability may result from a global decrease in GLT-1 expression and/or function. GLAST expression is restricted to glia, and glial cells provide upwards of four times the coverage of postsynaptic membranes compared to presynaptic membranes in the stratum radiatum (Lehre and Rusakov, 2002). As a result, GLAST may offer substantial postsynaptic protection when GLT-1 function is compromised. Alternatively, it is now well-established that approximately 5-10% of total GLT-1 protein is expressed in neurons, where it localizes to axon terminals themselves (Danbolt et al., 2016; Furness et al., 2008; Rimmele and Rosenberg, 2016; Zhou et al., 2019). Surprisingly, we have virtually no understanding of how this transporter pool regulates glutamate dynamics at the subcellular level. It is possible that this enigmatic presynaptic pool of GLT-1 is primarily affected in AD. Indeed, axonal GLT-1 is the main contributor to uptake in synaptosome preparations (Petr et al., 2015), and synaptosomal glutamate uptake is impaired in AD tissue (Hardy et al., 1987; Lauderback et al., 2001). Furthermore, neuronal GLT-1 knockout is sufficient to produce late-onset cognitive deficits (Sharma et al., 2019). By combining the imaging approach used in the current study with cell type specific knockouts of GLT-1, future studies can better understand how presynaptic GLT-1 shapes the subcellular profile of extracellular glutamate transients.

We used the DHK decay ratio as a measure of GLT-1’s contribution to total uptake at a given microenvironment; the greater the effect of saturating DHK being indicative of a greater role for GLT-1 in promoting rapid glutamate clearance (Hanson et al., 2015; Pinky et al., 2018). In agreement with a presynaptic vulnerability to glutamate spillover in AD, saturating DHK had a reduced effect at presynaptic membranes of 3xTg mice compared to WT controls.

Unexpectedly, the GLT-1 contribution to total glutamate uptake at the postsynaptic microenvironment was enhanced in 3xTg mice compared to WT controls. Thus, in 3xTg mice, GLT-1’s role in glutamate uptake is both impaired (presynaptic) and enhanced (postsynaptic) depending on the microenvironment in question. The precise mechanisms underlying this dichotomy remain to be addressed, but could represent a nanoscale malalignment of glial membranes, where the previously reported glial asymmetry favoring postsynaptic protection is exaggerated even further in 3xTg mice. Using super-resolution imaging techniques, it is of interest for future studies to better understand the nanoscale spatial relationships between GLT-1 expression, perisynaptic astrocytic processes, and pre- and postsynaptic membranes in 3xTg mice.

### Activity-dependent slowing of glutamate clearance

One notable feature of our iGluSnFR decay measures was that genotype differences were typically only detected after a longer train of neural activity was evoked with 100 pulses, and not with 5 pulses. The most likely explanation for this finding is suggested by the concept of spare glutamate transporters (Belo do Nascimento et al., 2021), and that the glutamate released by 5 pulses is insufficient to overwhelm the glutamate uptake system. In other words, 3xTg still have sufficient GLT-1 protein to efficiently clear glutamate released by short bursts of activity. In agreement with this explanation is the fact that ceftriaxone does not seem to accelerate glutamate clearance rates in WT animals (Wilkie et al., 2021). When glutamate transporters are challenged with HFS of longer duration (in this case, 1 second), uptake is slowed considerably. This activity-dependent slowing of glutamate clearance has been described previously (Armbruster et al., 2016; Pinky et al., 2018). During longer trains of neural activity, potassium efflux and electrogenic transporter currents depolarize astrocytic processes, which reduces the driving force for glutamate uptake (Armbruster et al., 2021). In the present study, it is possible that this natural activity-dependent slowing of glutamate clearance is exaggerated in 3xTg mice via an undefined mechanism. It will be of interest for future studies to test additional activity patterns, such as theta burst stimulation (which also causes activity-dependent slowing of glutamate clearance (Barnes et al., 2020), to determine the precise conditions under which 3xTg presynaptic membranes are exposed to excess glutamate.

### Differential regulation of glutamate dynamics at the subcellular level

In addition to their localization at pre- and postsynaptic membranes, glutamate receptors also exist on astrocytes. When we expressed iGluSnFR in astrocytes, we also observed slower uptake in 3xTg mice. Therefore, astrocytic glutamate receptors may also be vulnerable to overactivation in AD. Astrocytic NMDAR activation induces the release of pro-inflammatory cytokines, in particular TNF-alpha (Sühs et al., 2016). In addition to being a potent inflammatory cytokine, TNF-alpha is a powerful modulator of glutamate uptake, with TNF-alpha application to cultured astrocytes significantly reducing glutamate uptake (Ye and Sontheimer, 1996), at least in part through a downregulation of glutamate transporters (Zou and Crews, 2005). Importantly, TNF-alpha elevations are readily observed in the AD hippocampus (Jayaraman et al., 2021; Naghibi et al., 2021; Zhao et al., 2003). Therefore, it is possible that GLT-1 deficits promote the overactivation of astrocytic NMDARs, driving the release of TNF-alpha. It is important to consider that there is still much we do not know about astrocytic glutamate receptors-as significant heterogeneity exists in both the morphology and function of astrocytes, it is possible that the distribution of glutamate receptors on astrocytes may differ across brain regions (Emsley and Macklis, 2006; Höft et al., 2014; Regan et al., 2007; Sosunov et al., 2014). Indeed, astrocytic glutamate receptor activation has also been observed to confer neuroprotective roles (Durand et al., 2014; Jimenez-Blasco et al., 2015) in some studies.

Slow uptake in the 3xTg hippocampus was found to primarily affect presynaptic over postsynaptic membranes, and our subsequent electrophysiology experiments suggest that CA3-CA1 autoreceptor overactivation may act to oppose presynaptically-mediated post-tetanic potentiation in 3xTg mice. mGluRs are powerful modulators of synaptic transmission and their dysfunction is implicated in epilepsy, schizophrenia, and AD (Cosgrove et al., 2011). While mGluR expression is present throughout the hippocampus, mGluR5 and mGluR7 have been detected in CA3 in high concentrations, and their activation leads to a depression of CA3-CA1 synaptic neurotransmission (Gereau IV and Conn, 1995; He et al., 2019; Klar et al., 2015; Pittaluga, 2016; Shigemoto et al., 1997). Our electrophysiology experiments support this observation, as we found that PTP is decreased in 3xTg mice. Moreover, mGluR5 (MTEP) or mGluR7 (MSOP) antagonism restored PTP values to WT levels. Interestingly, ceftriaxone alone was sufficient to restored PTP to control levels, suggesting that the glutamate accumulation at 3xTg axons can overstimulate mGluR autoreceptors to oppose plasticity. It is well-known that synapse loss is one of the best correlates of cognitive decline in AD (Colom-Cadena et al., 2020), and synapse loss in the hippocampus is reported in both preclinical and clinical studies of AD (Leshchyns’ka et al., 2015; Neuman et al., 2015). Interestingly, ceftriaxone administration improves cognitive function in 3xTg mice (Zumkehr et al., 2015); thus, an intriguing possibility is that an early presynaptic vulnerability to glutamate spillover may promote subsequent synapse elimination and cognitive decline in AD.

### Summary and conclusions

By isolating the study of extracellular glutamate dynamics at pre- or postsynaptic microenvironments, we have revealed a functional consequence of the asymmetric morphological arrangement of astrocytes first described almost two decades ago by Lehre and Rusakov (2002). Despite an often postsynaptic-centric view of the consequences of glutamate spillover in disease, we reveal a presynaptic vulnerability to GLT-1 impairment in the 3xTg model of AD that opposes short-term potentiation at CA3-CA1 synapses. Our results shed new light on the consequences of GLT-1 dysfunction in AD and may have broader implications for presynaptic vulnerability in a range of disease states associated with GLT-1 reduction.

