## Supplemental Figures 1-8 for "Asymmetric dysregulation of glutamate dynamics across the synaptic cleft in a mouse model of Alzheimer disease"

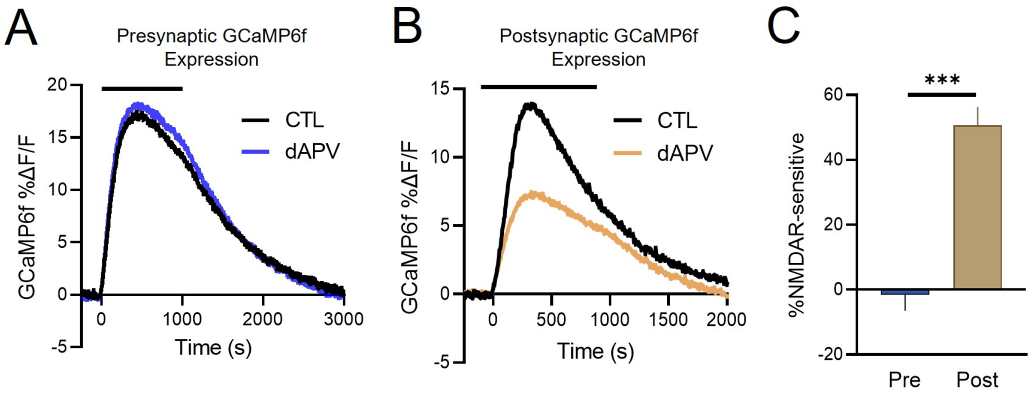


**Supplementary Figure 1. NMDA receptor blockade reduces postsynaptic but not presynaptic calcium responses to high-frequency stimulation.** (A) Presynaptic GCaMP6f response to high-frequency stimulation (HFS) before (black) and after (blue) bath application of d-APV (50 µM). (B) Postsynaptic GCaMP6f response to HFS before (black) and after (orange) d-APV. (C) postsynaptic GCaMP responses are more sensitive to NMDAR blockade than presynaptic GCaMP responses, with a 50% reduction observed in the postsynaptic GCaMP response. Error bars represent s.e.m. *** $p < 0.001$.

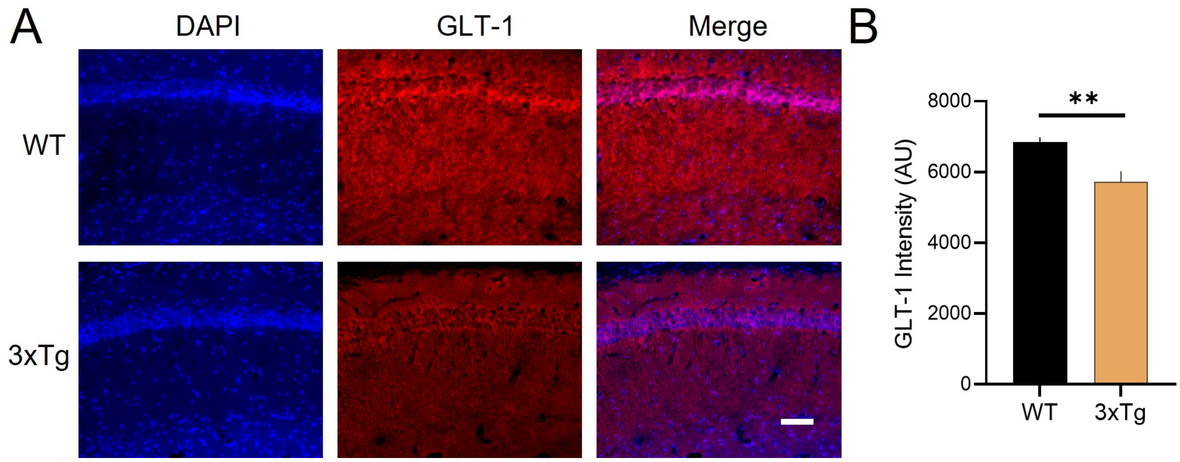


**Supplementary Figure 2: GLT-1 expression is significantly reduced in 3xTg hippocampus.** WT and 3xTg mice were perfused at 6 months of age. GLT-1 intensity was quantified in stratum radiatum. All immunostaining was performed at the same time and imaging parameters (LED intensity, exposure times) remained consistent for both genotypes. WT n = 12, 3xTg n = 10. Scale bar in A: 50 µm. Error bars represent s.e.m. ** $p < 0.01$.

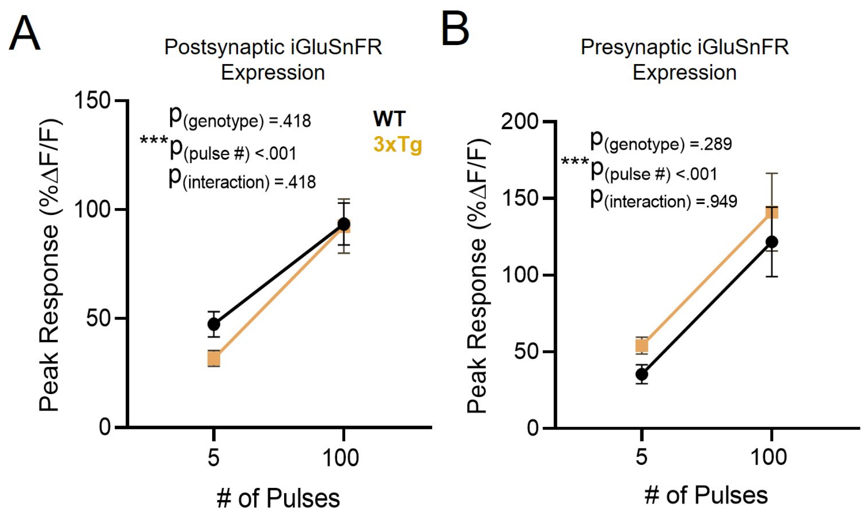


**Supplementary Figure 3. Peak iGluSnFR responses do not differ between WT and 3xTg mice.** (A) Postsynaptic iGluSnFR response peaks in WT (black) and 3xTg (orange) mice. (B) Presynaptic iGluSnFR response peaks in WT (black) and 3xTg (orange) mice.

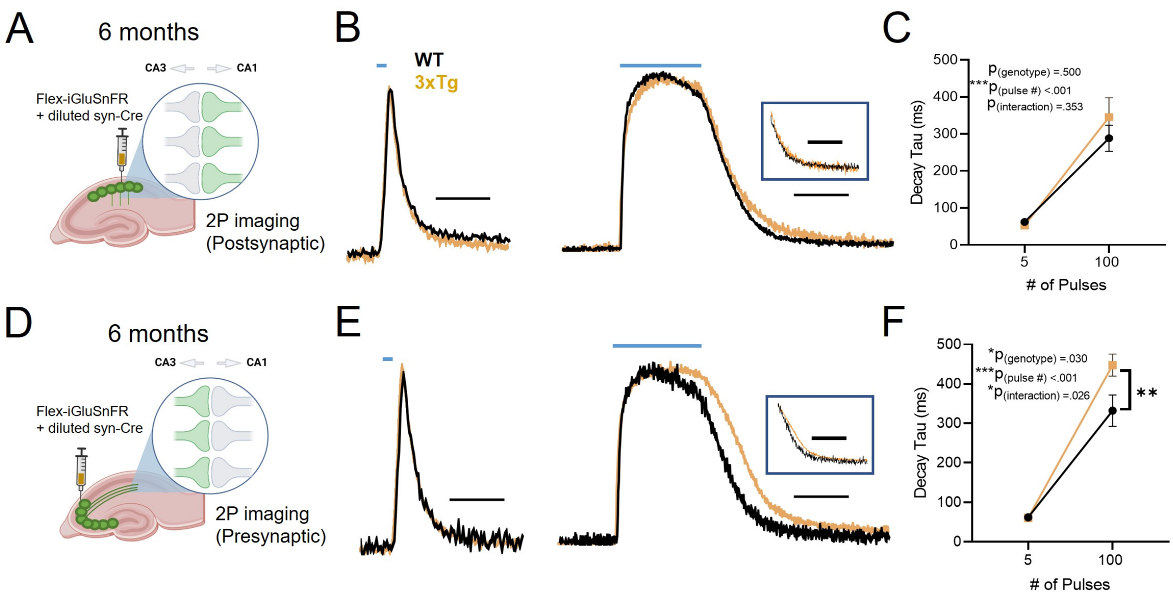


**Supplementary Figure 4. Presynaptic glutamate clearance impairment and spared postsynaptic clearance in the 3xTg hippocampus replicated at 32 degrees.** (A-C) Postsynaptic iGluSnFR expression (A). Average traces in WT (black) and 3xTg mice (orange) in response to 5 (B, left) or 100 (B, right) pulses of evoked activity. Grouped data are shown in C. (D-F) Same as A-C but for presynaptic iGluSnFR expression. All experiments conducted in ACSF heated to 32 °C. Horizontal blue lines indicate the timing and duration of electrical stimulation. Scale bars in B and E: 200 ms (left) and 500 ms (right). Traces in boxes show average iGluSnFR responses normalized to the value at the end of the one second of electrical stimulation. Error bars represent s.e.m. $*p < 0.05$, $***p < 0.001$.

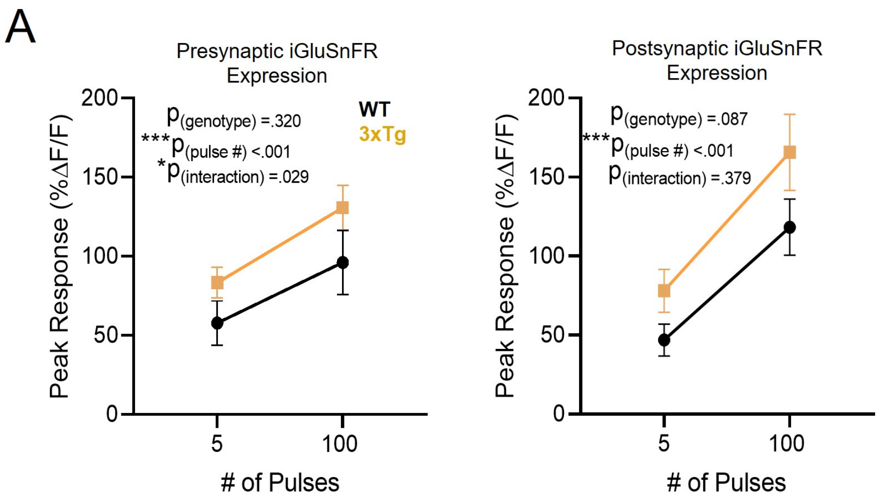


**Supplementary Figure 5. Peak iGluSnFR responses do not differ between WT and 3xTg mice after GLT-1 blockade with DHK.** (A) Presynaptic iGluSnFR response peaks in WT (black) and 3xTg (orange) mice. (B) Postsynaptic iGluSnFR response peaks in WT (black) and 3xTg (orange) mice. Response peaks were obtained in the presence of a saturating concentration (300 µM) of the GLT-1 inhibitor DHK.

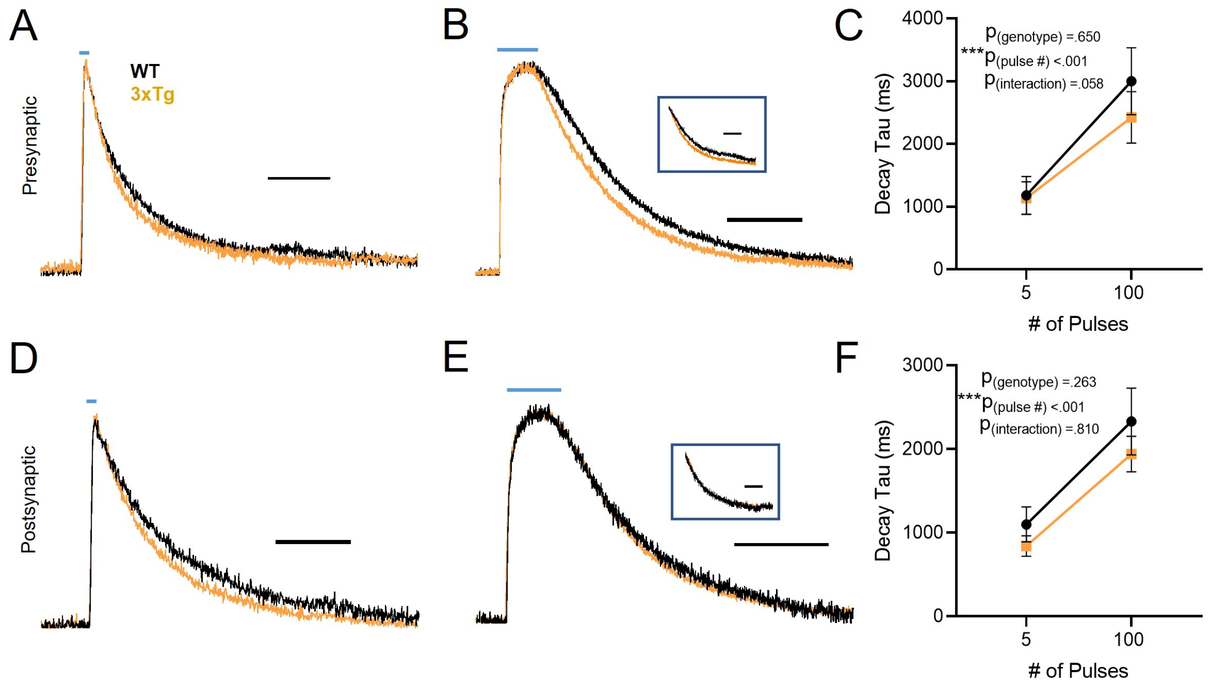


**Supplementary Figure 6. Diffusion does not differ between WT and 3xTg mice.** (A-C) Presynaptic iGluSnFR responses to 5 (A) and 100 (B) pulses in the presence of 100 µM TBOA to block transporter-mediated uptake. (D-F) Same for A-C but for postsynaptic iGluSnFR expression. Horizontal blue lines indicate the timing and duration of electrical stimulation. Scale bars in A and D:  1000 ms. Scale bars in B and E: 2000 ms. Traces in boxes show average iGluSnFR responses normalized to the value at the end of the one second of electrical stimulation. Error bars represent s.e.m.

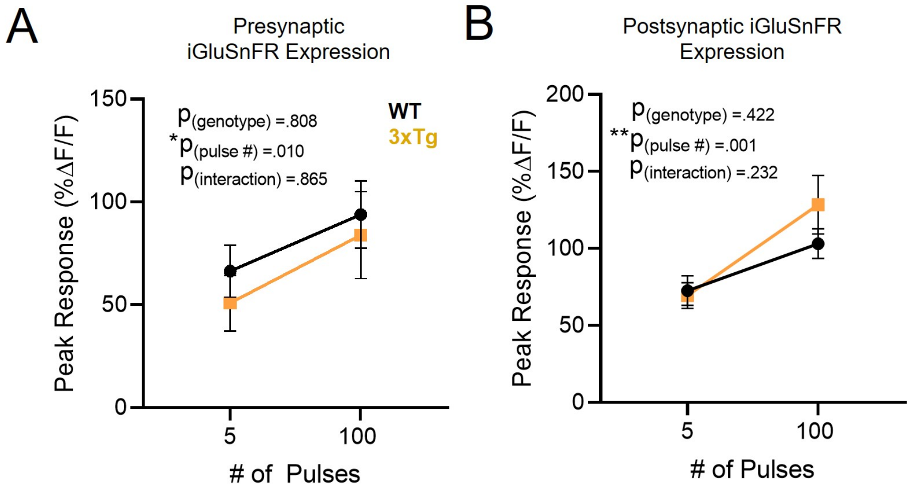


**Supplementary Figure 7. Peak iGluSnFR responses do not differ between WT and 3xTg mice after non-selective glutamate transporter blockade with TBOA.** (A) Presynaptic iGluSnFR response peaks in WT (black) and 3xTg (orange) mice. (B) Postsynaptic iGluSnFR response peaks in WT (black) and 3xTg (orange) mice. Response peaks were obtained in the presence of a saturating concentration (300 µM) of the GLT-1 inhibitor DHK.

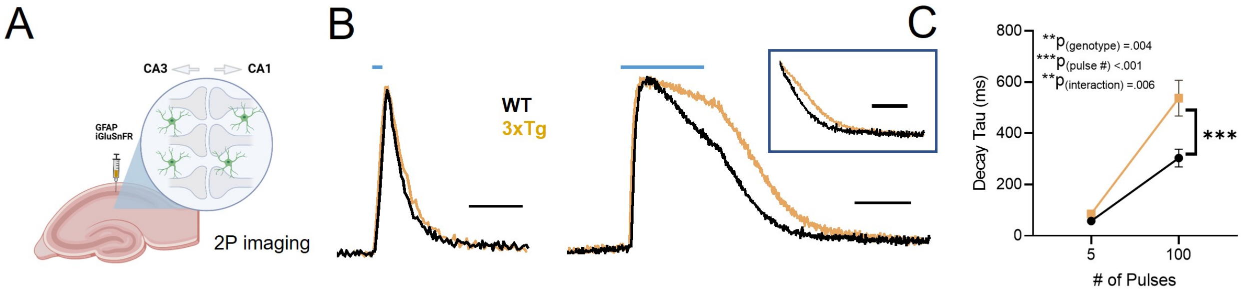


**Supplementary Figure 8. Glutamate clearance is significantly slower in astrocytes of the 3xTg hippocampus.** (A) Schematic of GFAP-iGluSnFR. (B) Average iGluSnFR responses to 5 (left) and 100 (right) pulses of stimulation in WT (black) and 3xTg (orange) mice. Grouped data shown in (C). Horizontal blue lines indicate the timing and duration of electrical stimulation. Scale bar in B: 200 ms. Scale bar in C: 500 ms. Traces in box show average iGluSnFR responses normalized to the value at the end of the one second of electrical stimulation. Error bars represent s.e.m. *** $p < 0.001$.
